# Brain-wide functional networks associated with anatomically- and functionally-defined hippocampal subfields using ultrahigh-resolution fMRI

**DOI:** 10.1101/2020.12.01.407049

**Authors:** Wei-Tang Chang, Stephanie K. Langella, Yichuan Tang, Han Zhang, Pew-Thian Yap, Kelly S. Giovanello, Weili Lin

**Author notes:** Corresponding authors at: **Biomedical Research Imaging Center,** University of North Carolina at Chapel Hill, NC, USA. E-mail address (W.-T. Chang) and (W. Lin).

## Abstract

The hippocampus is critical for learning and memory and may be separated into anatomically-defined hippocampal subfields (aHPSFs). Many studies have shown that aHPSFs, and their respective functional networks, are differentially vulnerable to a variety of disorders. Hippocampal functional networks, particularly during resting state, are generally analyzed using aHPSFs as seed regions, with the underlying assumption that the function within a subfield is homogeneous, yet heterogeneous between subfields. However, several prior studies that have utilized aHPSFs and assessed brain-wide cortical connectivity have observed similar resting-state functional connectivity profiles between aHPSFs. Alternatively, data-driven approaches offer a means to investigate hippocampal functional organization without a priori assumptions. However, insufficient spatial resolution may lead to partial volume effects at the boundaries of hippocampal subfields, resulting in a number of caveats concerning the reliability of the results. Hence, we developed a functional Magnetic Resonance Imaging (fMRI) sequence on a 7T MR scanner achieving 0.94 mm isotropic resolution with a TR of 2s and brain-wide coverage to 1) investigate whether hippocampal functional segmentation with ultrahigh-resolution data demonstrate similar anatomical, lamellar structures in the hippocampus, and 2) define and compare the brain-wide FC associated with fine-grained aHPSFs and functionally-defined hippocampal subfields (fHPSFs). Using a spatially restricted hippocampal Independent Component Analysis (ICA) approach, this study showed that fHPSFs were arranged along the longitudinal axis of the hippocampus that were not comparable to the lamellar structures of aHPSFs. Contrary to the anatomically defined hippocampal subfields which are bilaterally symmetrical, 13 out of 20 fHPSFs were unilateral. For brain-wide FC, the fHPSFs rather than aHPSFs revealed that a number of fHPSFs connected specifically with some of the functional networks. The visual and sensorimotor networks preferentially connected with different portions of CA1, CA3 and CA4/DG. The DMN was also found to connect more extensively with posterior subfields rather than anterior subfields. Finally, the frontoparietal network (FPN) was anticorrelated with the head portion of CA1. The investigation of functional networks associated with the fHPSFs may enhance the sensitivity of biomarkers for a range of neurological disorders, as network-based approaches take into account disease-related alterations in brain-wide interconnections rather than measuring the regional changes of hippocampus.

## INTRODUCTION

The hippocampus plays a major role in learning and memory (Squire et al., 1992, Tulving and Markowitsch, 1998, Spaniol et al., 2009). Cytoarchitectural, connectomic, and neurophysiological studies have revealed that the hippocampus is a heterogeneous neural structure with complex efferent and afferent connections with multiple brain cortices. Therefore, it is not surprising that extensive efforts have been devoted to defining hippocampal subfields as well as constructing brain-wide hippocampal-subfield specific functional networks. In particular, two different approaches have been widely used to determine hippocampal subfields, namely anatomically and functionally defined subfields. Anatomically, the hippocampal formation consists of multiple, lamellar subdivisions organized along the longitudinal axis of hippocampus, including cornu ammonis 1 (CA1), CA3, CA4, dentate gyrus (DG) and subiculum (Duvernoy et al., 2013, Ding et al., 2016) (Andersen et al., 1971). Hippocampal subfields have been reported to be differentially vulnerable to a variety of disorders, including autism, Alzheimer’s disease, schizophrenia, and bipolar disorder (Mueller et al., 2008, Neylan et al., 2010, La Joie et al., 2013, Schoene-Bake et al., 2014, de Flores et al., 2015, Hsu et al., 2015), as well as prodromal disease states and in individuals clinically normal, yet at-risk for disease (Dournal, et a;., 2020; McKeever, et al., 2020; Nadal et a;, 2020; Nurdal et al., 2020).

In contrast, recent investigations, including genetic, as well as functional studies using electrophysiological or fMRI techniques, have revealed that the hippocampal functions are regionally specialized. Using data-driven approaches examining the blood-oxygen-level-dependent (BOLD) signal characteristics of the entire hippocampus, e.g., independent component analysis (ICA), prior studies have reported that functionally independent hippocampal components were spatially arranged along the longitudinal axis (Blessing et al., 2016), as well as medial–lateral axis (Plachti et al., 2019). Similar findings were also reported by Zhong et al. who utilized functional correlations between hippocampal voxels and whole-brain cortical regions (Zhong et al., 2019). Together these studies appear to suggest that the general feature of hippocampal subfields is independent of the approaches (anatomy or functions) used to defined them. That is, the hippocampal subfields are organized along the longitudinal axis as well as potentially medial-lateral axis. Despite this consistent general feature, discrepancies exist using the two approaches to define hippocampal subfields. First, there are distinct anatomical demarcations between anatomically and functionally defined subfields. The anatomically-defined hippocampal subfields have a lamellar structure and the functionally-defined hippocampal subfields exhibit a barrel-shaped structure (Fanselow and Dong, 2010, Poppenk et al., 2013, Strange et al., 2014). Second, the brain-wide networks defined using the two approaches differ. Specifically, one of the main assumptions of using anatomically defined subfields as seeds to construct brain-wide functional networks is that within-subfield functional characteristic is homogenous, while across-subfield functional characteristic is heterogeneous. Such an assumption, however, have been challenged by several studies, reporting that resting-state activities between hippocampal subfields are highly correlated (Lacy and Stark, 2012, Shah et al., 2018) and that the functional networks constructed using anatomically defined subfields highly resembles that obtained using the entire hippocampus as a seed region (Bai et al., 2011, Wang et al., 2015). In contrast, using functionally defined subfields, the brain-wide networks associated with anterior and posterior hippocampus exhibit distinct patterns (Blessing et al., 2016). Furthermore, Zhong et al. operatively subdivided the hippocampus into head, body and tail portions and showed that the head portion of hippocampus demonstrated a higher connectivity strength with the sensorimotor network than that of body and tail portions (Zhong et al., 2019).

A caveat to the above findings, however, is that most of the prior hippocampal-subfield studies employed relatively low spatial resolutions (≥ 2 mm isotropic) (Zarei et al., 2013, Blessing et al., 2016, de Flores et al., 2017, Vos de Wael et al., 2018, Zhong et al., 2019) and/or an insufficient anatomical coverage. Insufficient spatial resolution may not only lead to partial volume effects at subfield boundaries, but also reduce the effective size of unmixed voxels. As a result, the mixture of subfield signals could result in false positive FC as well as affect the functional parcellation of hippocampus (Plachti et al., 2019, Zhong et al., 2019). To mitigate the shortcomings attributed by the limited spatial resolution, several approaches capable of acquiring ultra-high spatial resolution fMRI have been proposed. However, these approaches compromise either the ability to acquire whole brain images (Libby et al., 2012, Dalton et al., 2019b) and/or to achieve a high temporal resolution (>3 s) (Kahn et al., 2008, Bergmann et al., 2016). A limited spatial coverage makes it impractical to assess brain-wide hippocampal subfield functional networks whereas low temporal resolution renders the images vulnerable to signal contamination from other physiological signals (cardiac and/or respiratory signals).

To this end, we developed a fMRI sequence achieving 0.94 mm isotropic resolution, a repetition time (TR) of 2 s, and whole-cerebrum coverage on 7T, mitigating the aforementioned limitations (Chang and Lin, 2018) (See Supplementary materials for the details of the imaging methods). Using the newly developed approach, this study aimed to 1) address whether hippocampal functional subfields are similar to anatomically-defined lamellar structures and 2) define and compare the brain-wide FC associated with fine-grained aHPSFs and fHPSFs using ultrahigh-resolution fMRI images. To address the first aim, segmentation was performed using ICA and k-means clustering based on the BOLD time-series of hippocampal voxels. For the second aim, the iFC maps associated with aHPSFs and fHPSFs as well as the spatial distribution of iFC maps in relation to the resting-state networks (RSNs) were demonstrated. A quantitative measure was developed to quantify the functional-network specificity for the aHPSF or fHPSF. To understand how different hippocampal segmentations affect the detection and strength of cotical-hippocampal subfield FC, the hippocampal coverages of significant iFC by every cortical vertex were demonstrated. The results associated with aHPSF and fHSPF were compared..

## MATERIALS AND METHODS

### Participants

Sixteen healthy participants (aged 27.9±7.2 years, 8 females) were recruited for this study. Participants were screened to ensure they have no history of neurological or psychiatric conditions, previous head trauma, or MRI contraindications. Informed consent was obtained from each participant prior to the study in accordance with the experimental protocol approved by the local Institutional Review Board.

### MR Acquisitions

MR images were acquired using a 7T Magnetom scanner (Siemens Healthcare, Erlangen, Germany). MPRAGE images were acquired first using the following imaging parameters: TR/TE/TI = 2200/2.79/1050 ms, flip angle = 7°, partition thickness = 0.94 mm, image matrix = 256 × 208, 192 partitions, and FOV = 24.0 cm × 19.5 cm. Subsequently, two 6-minute resting-state scans were acquired using the blipped Partition-encoded Simultaneous Multi-Slab (bPRISM) sequence. The details of the bPRISM sequence can be found in the Supplementary Materials. The participants were instructed to remain awake with eye opened during the resting-state scans. The spatial resolution was 0.94 mm isotropic using the following imaging parameters: FOV = 169×112.7×164.5 (R-L×H-F×A-P) mm^3^; SMS factor = 7; TE = 23 ms; TR per slab volume = 400 ms; effective TR = 2 s; and number of repetitions 180.

### Data Preprocessing

Time-series data were motion corrected using FSL (Smith et al., 2004). The motion-corrected data were decomposed into a number of independent components by MELODIC (Beckmann et al., 2005). With the 32 resting-state runs acquired from 16 participants, we trained the ICA-based denoising tool FIX to auto-classify ICA components into signal and noise components (Salimi-Khorshidi et al., 2014). The noise components were then removed by FIX using a threshold of 20. The time-series signals were band-pass filtered from 0.01 to 0.1 Hz.

### Spatial Co-registrations

To minimize crosstalk between hippocampal voxels, the analysis of functional connectivity (FC) was performed on the individual subject’s native EPI space in order to avoid spatial blurring induced by either distortion correction or co-registration. The individual T1 images were coregistered onto the EPI using ANTS (Advanced Normalization Tools 2.1.0) (Avants et al., 2011) as shown in Figure 1. To improve coregistration quality, exterior CSF signals of the EPI images were automatically labeled using FSL FAST (Zhang et al., 2001) and removed. The T1 images were also masked by the union of gray matter and white matter in order to remove interference from the brain meninges. Nonlinear coregistration with the similarity metric of probability mapping was used.

**Figure 1:**
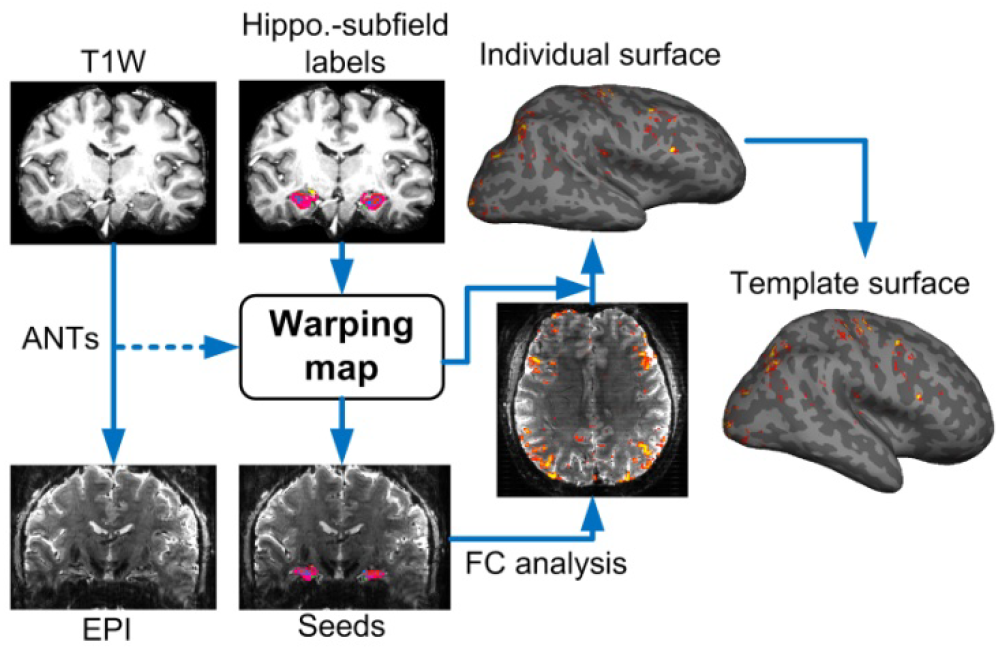
The procedures of coregistration. Individual T1-weighted images were coregistered to the EPI by ANTs. The warping coefficients were then applied to the labeled images of the hippocampal subfields. The seed-based FC analysis was performed on the EPI space and the FC map was re-sampled onto individual cortical surface before being warped to the template surface. The re-sampling of FC values on EPI space captured only the signal from the white matter surface to the middle depth of cerebral cortex in order to suppress contamination from large vessels.

### Anatomical Segmentation of Hippocampus

The labels of aHPSFs were obtained using Freesurfer 6.0 (Fischl, 2012, Iglesias et al., 2015, Saygin et al., 2017) on T1 images (Figure 1). This study employed the version released on August 31^st^, 2017 which is able to subdivide the hippocampal substructures into head, body and tail where applicable. Specifically, the presubiculum, subiculum, CA1, CA3, CA4/DG are subdivided into head and body portions. Together with parasubiculum and hippocampal tail, we identified 24 aHPSFs (12 per hemisphere) for the following analyses. After coregistering the individual T1 images onto the EPI images, seed-based functional connectivity of each subfield was calculated on EPI space before being warped back to the T1 images.

### Functional Segmentation of Hippocampus

Spatially-restricted ICA and k-means clustering were used to perform functional segmentation of the hippocampus (see Figure S6). The EPI time-series signals of each individual were coregistered onto the template brain by resampling the image-series once using the concatenated transformations, which included the transformations from individual EPI, T1 to template spaces. The transformation from individual EPI to T1 images was the inverse of the transformation from individual T1 images to EPI as shown in Figure 1. The transformations from individual T1 to template brains were estimated using the masked hippocampus instead of the whole brain. This step is needed owing to the fact that the cortical folding pattern of each brain varies considerably, leading to inaccurate hippocampal coregistration if the whole brain transformation is used between individual brain images and the template.

Subsequently, the hippocampal temporal signals of all subjects were warped onto the hippocampal template. The group ICA was performed using MELODIC (Beckmann et al., 2005). The number of independent components for MELODIC operation was set as 24, which is the total number of aHPSFs. The outputs of MELODIC included the probability maps of the independent components. The k-means clustering method utilized the probability maps as the feature vector and generated the segmentation maps. Although the cluster number was initially set as 24, the final cluster number was determined automatically according to the elbow criterion of the cluster validity index, which is defined as the ratio of within-cluster distances to between-cluster distances (Shen et al., 2016).

### Statistical Mapping of Functional Connectivity on the Cortical Surface

The time course associated with each seed was obtained by averaging all time courses within each of the aHPSF or fHPSF. To suppress the signal contamination from pial vessels, only the values from the middle depth of the cortex to the gray-white matter boundary were sampled and averaged. Surface-based spatial smoothing was then applied with full-width half-magnitude (FWHM) of 3 mm. This method has been shown to improve inter-subject consistency on functional mapping in our previous work (Ahveninen et al., 2016). After the intrinsic-FC (iFC) values were warped from individual EPI space onto the cortical surface of the template brain, statistical significance of the group-averaged Fisher’s z values was calculated by a t-test using the Freesurfer command ‘mri_glmfit’. In order to minimize the effect of spatially-varied signal loss and gender effects, the averaged tSNR of the seed region and gender were incorporated into the group analysis as covariates. Correcting for multiple comparisons was performed by the Monte Carlo Simulation in FreeSurfer. The number of permutations was set to 4000. The vertex-wise and cluster-wise p-value thresholds were 0.05. The corrected p-values were converted back to z scores for better visualization.

### Measure of Functional-Network Specificity

To better understand how the spatial distribution of iFC maps in relation to commonly defined brain functional networks, seven RSNs using the functional atlas reported by Yeo et al (Yeo et al., 2011) were employed. In this study, we developed a measure called non-uniformity to evaluate whether a hippocampal subregion (aHPSF or fHPSF) was connected mainly with some specific functional networks or uniformly with many functional networks. Here we defined c_i_ as the percentage of the *i*^th^ functional network that was significantly connected with a hippocampal subregion and **c** is the array of the coverage percentages. To test whether a hippocampal subregion connected to the functional networks uniformly or selectively, we defined the non-uniformity ν as

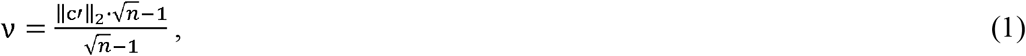

where **c**′ = **c**/Σ_*i*_ *c*_*i*_ and *n* denotes the total number of functional networks. If a hippocampal subregion connected to all the functional networks with the same coverage, the ||**c**′||_2_ becomes 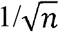 and the non-uniformity ν becomes 0. If a hippocampal subregion connected to only one functional network, on the other hand, the ||**c**′||_2_ is 1 and the ν is 1. Hence, ν ranges from 0 to 1 where 0 represents complete uniformity and 1 represents the highest non-uniformity, i.e. highest specificity. High specificity in this study implied that a hippocampal subregion connected with the functional networks selectively.

## RESULTS

The surface reconstructions of aHPSFs and fHPSFs are shown in Figure 2a and 2b respectively. A total of 20 fHPSFs were identified. To demonstrate the spatial relationship between aHPSFs and fHPSFs, the overlapped percentages of each fHPSF with every aHPSF are shown in Figure 2b In general, aHPSFs spatially extended along the longitudinal axis of hippocampus, while fHPSFs consisted of barrel-shaped components primarily along the longitudinal axis, consistent with many reports of longitudinal functional specialization (Moser and Moser, 1998, Fanselow and Dong, 2010, Poppenk et al., 2013, Strange et al., 2014). No functionally-defined hippocampal subfields demonstrate lamellar structures similar to aHPSFs. Contrary to the anatomically defined hippocampal subfields which are bilaterally symmetrical, 13 out of 20 fHPSFs were unilateral, suggesting that the iFC of hippocampus is not symmetrical between the two hemispheres. For example, the tail of hippocampus is separated into two functional components, fHPSF #15 and #20, for right and left of hippocampus, respectively. Furthermore, the fHPSF #3 spatially spanned across head, body and tail of hippocampus and overlaped with 56% and 41% of the head and body portion of left CA1, respectively..

**Figure 2:**
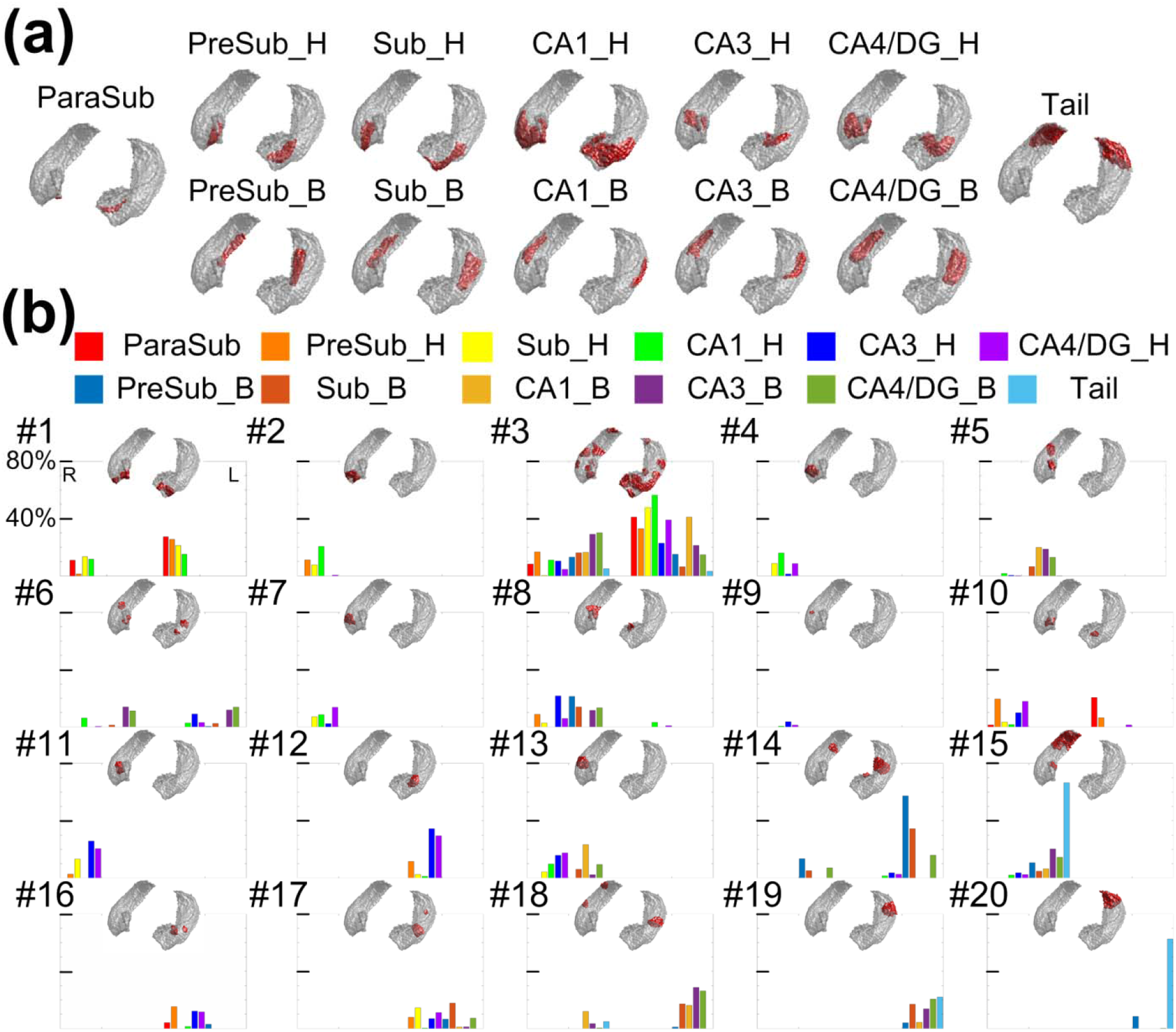
Anatomically and functionally defined hippocampal subfields and their sptaial relationships. (a) Surface reconstruction of aHPSF. (b) Surface reconstruction of fHPSF and the percent of spatial overlap between aHPSF and fHPSF. The overlapped percentages with different aHPSFs are color-encoded as indicated in the figure legend. The indices of fHPSFs were arranged from anterior to posterior. The index #1 was assigned to the most anterior fHPSF and #20 to the most posterior fHPSF.

The iFC maps associated with aHPSFs and fHPSFs are shown in Figure 3a and 3b respectively. The first two maps from the left of each row represent the lateral view of cortical surface and the following two maps represent the medial view of cortical surface. The maps in the left and right sides of Figure 3a are associated with the aHPSFs in the left and right hemispheres respectively. The z scores of FC are color-encoded as indicated by the color bar. The merged iFC maps across aHPSFs and fHPSFs covered 67.3% and 64.3% of cortical surface respectively. Additionally, the overlapping percentages of merged iFC maps with each of the resting-state functional networks are listed in Table 1. The overlapping ratio between the two merged iFC maps was 72.5%, indicating that the aHPSFs and fHPSFs have similar topologies of the merged iFC maps. Hence, the corresponding analyses of hippocampal subfields are minimally biased by the detection sensitivity difference of iFC between aHPSF and fHPSF.

**Figure 3:**
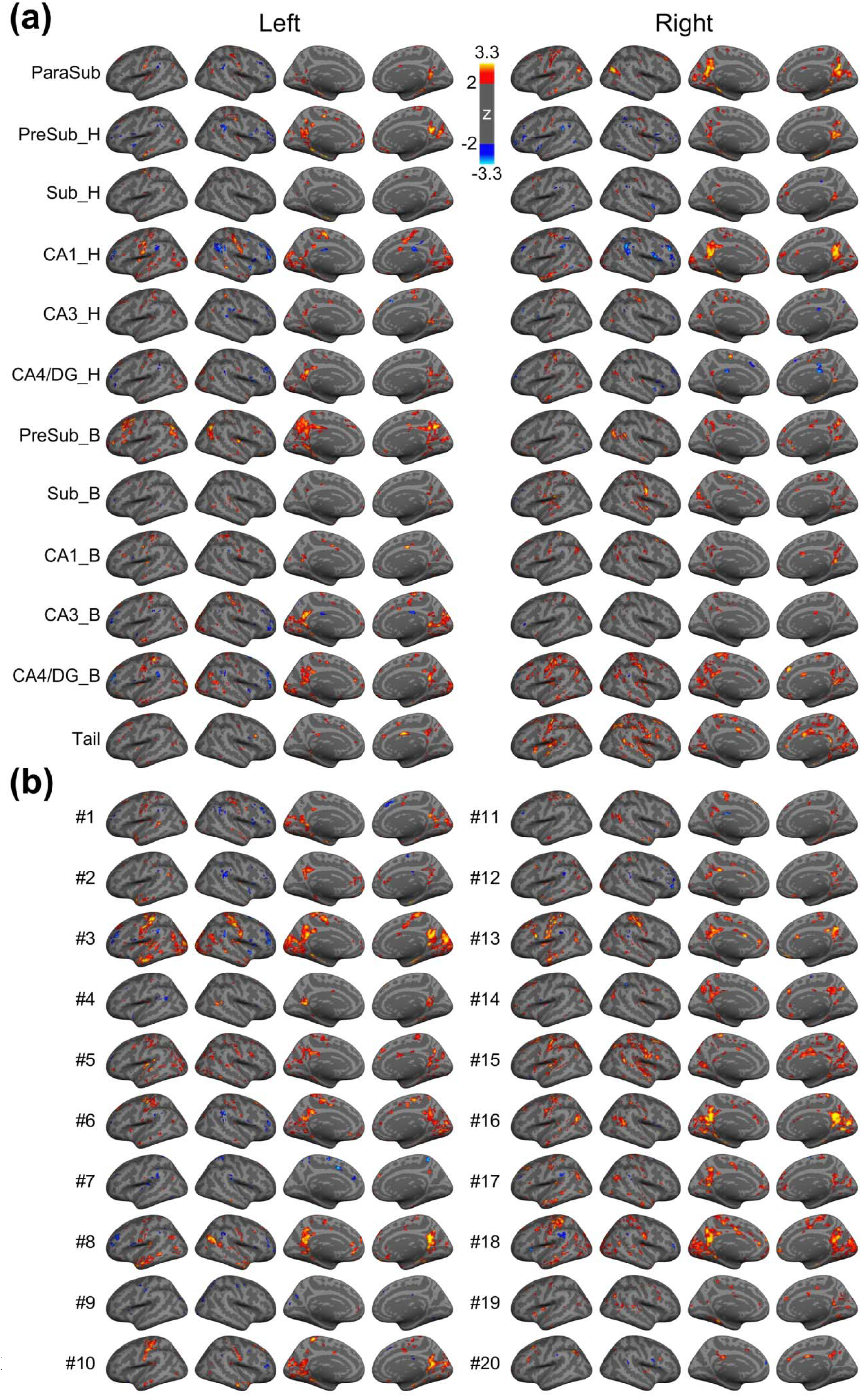
Whole brain functional connectivity maps using (a) aHPSFs and (b) fHPSFs as seed regions, respectively. The statistical significances of iFC maps were assessed using a vertex-wise non-parametric permutation test with 4000 permutations. The statistical maps of iFC were corrected for multiple comparisons and cluster-wise threshold set to p < 0.05. The z scores are color-encoded as indicated by the color bar. Abbreviations: H – head; B – body; Sub – subiculum; CA – cornu ammonis; DG – dentate gyrus.

**Table 1:**
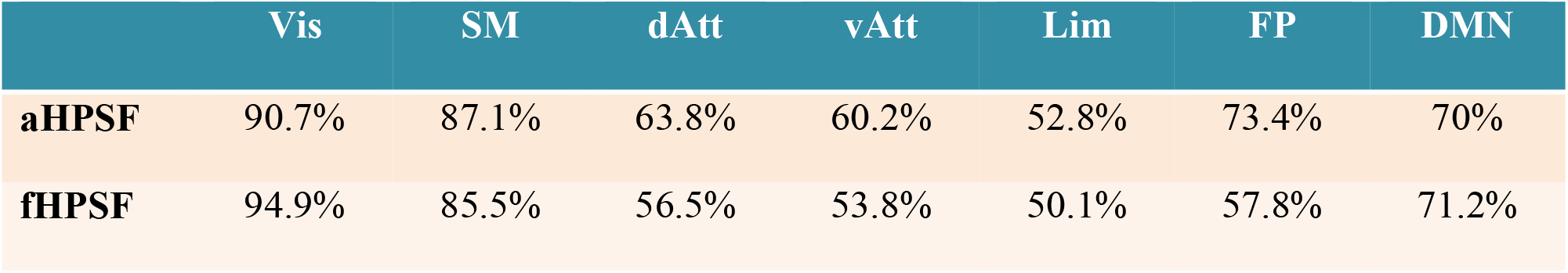
The overlapping percentages of merged iFC maps with different resting-state functional networks. Abbreviations are as described in Figure 3.

|  | Vis | SM | dAtt | vAtt | Lim | FP | DMN |
| --- | --- | --- | --- | --- | --- | --- | --- |
| **aHPSF** | 90.7% | 87.1% | 63.8% | 60.2% | 52.8% | 73.4% | 70% |
| **fHPSF** | 94.9% | 85.5% | 56.5% | 53.8% | 50.1% | 57.8% | 71.2% |

To demonstrate the spatial distribution of iFC maps in relation to the commonly defined brain networks, the percentages of the iFC maps with respect to the resting-state functional networks are shown by radar charts in Figure 4. The radar charts of aHPSFs and fHPSFs were sorted together based on the functional-network specificity (see Materials and Methods) and divided into three groups with the high 1/3, medium 1/3 and lowest 1/3 of specificity. In the high-specificity group, 10 out of 15 charts are of fHPSF even though the total number of fHPSF is lower than that of aHPSF. These results suggest that the fHPSF provides higher sensitivity than aHPSF to detect the iFC specificity to functional networks. Furthermore, for each fHPSF in the group of high specificity, the functional networks that received the highest coverages of iFC are visual, sensorimotor or default-mode networks. The highest iFC coverage of visual network is 67% while those of sensorimotor and default-mode networks are 48% and 27% respectively. The FC coverages of the other functional networks are all below 16%, suggesting that the high functional-network specificity is associated more with visual, sensorimotor and default-mode networks than other networks in resting state.

**Figure 4:**
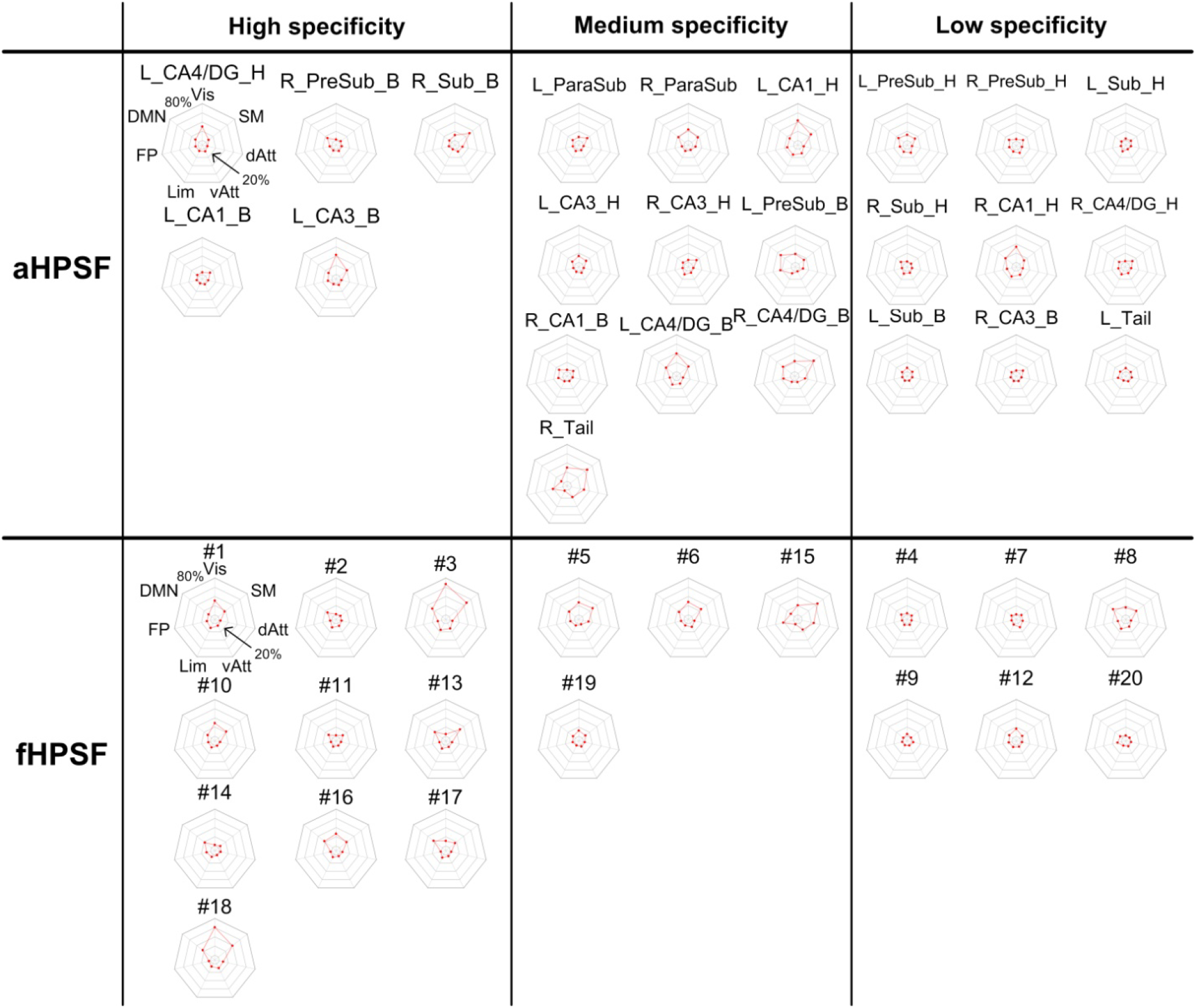
The percentages of iFC coverage of each RSN associated with aHPSFs and fHPSFs. The radar plots in the right-most columns of each panel shows the percentages of each iFC map with respect to the seven intrinsic functional networks. The innermost and outermost isoparametric curves in the radar plots represent 0% and 80% respectively. The increment between adjacent isoparametric curves is 20%. The charts of aHPSFs and fHPSFs are sorted together based on the hippocampal specificity and then classified into three groups. Abbreviations: L – left; R – right; other abbreviations are as described in Figure 3 and Table 1.

For better visualization, we collated the information from all the charts in Figure 4 and reorganized in Figure 5. Only the RSNs with the 4 highest coverage of iFC are shown. The head portions of CA1 connect extensively with visual and sensorimotor networks (maximum at 42% and 20%, respectively) but the body portions show much less cortical connections (maximum at 12%). On the contrary, the body portions of CA3 and CA4/DG show more cortical connections with visual and sensorimotor networks (maximum at 42% and 43% respectively) than the head portions (maximum at 13%). Among all the aHPSFs, the body portion of pre-subiculum has most extensive FC with DMN (31%).

**Figure 5:**
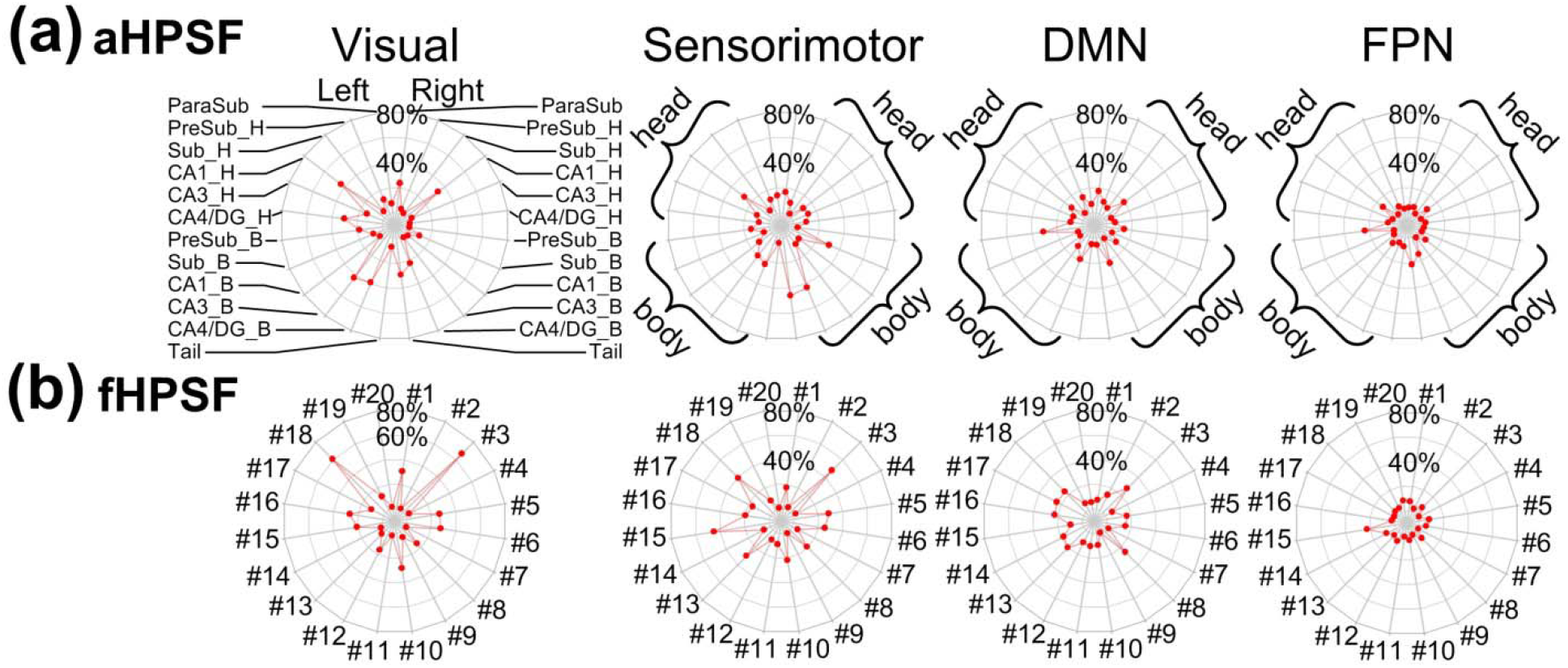
The percentages of iFC coverage of each RSN associated with (a) aHPSFs and (b) fHPSFs. The radar charts from left to right panels correspond to visual network, sensorimotor network, DMN and FPN. The names of aHPSF are only illustrated in the left most panel in (a). The head and body portions of aHPSF are highlighted by curly brackets in the other panels. For the radar charts of fHPSF in (b), the indices of fHPSFs were arranged from anterior to posterior. The index #1 was assigned to the most anterior fHPSF and #20 to the most posterior fHPSF. The innermost and outermost isoparametric curves in the radar plots represent 0% and 80% respectively. The increment between adjacent isoparametric curves is 20%. Abbreviations: L – left; R – right; H – head; B – body; Sub – subiculum; CA – cornu ammonis; DG – dentate gyrus; FPN – frontoparietal network; DMN – default-mode network.

For the iFC of fHPSFs, as shown in Figure 3b and 4b, the fHPSF #3 consists of head, body and tail portions and has the most extensive FC with visual network, sensorimotor network and DMN (67%, 48% and 27% respectively). The FC patterns of fHPSF #3 and the head portion of left CA1 are similar (correlation coefficient = 0.49, *p* < 0.001), which reflect the fact that fHPSF #3 overlaps with 56% of the CA1 head and 41% of the CA1 body. From the RSN’s perspective, the visual network shows extensive FC with fHPSF #1 (31%), #5 (27%), #6 (28%), #10 (28%), #16 (26%) and #18 (61%). Likewise, the sensorimotor network has extensive FC with fHPSF #5 (29%), #6 (26%), #13 (30%), #15 (45%) and #18 (39%). These results show that the visual and sensorimotor networks connect with distributed anatomical regions along the hippocampal longitudinal axis but some fHPSFs in the posterior hippocampus are more prominent. DMN shows the FC primarily with fHPSF #3 (27%), #8 (25%), #13 (20%), #16 (22%), #17 (24%) and #18 (24%). Almost all of the corresponding fHPSFs are either closed to or in the body portion but not the head portion of hippocampus. Different from the visual network, sensorimotor network and DMN, the FPN has much less FC with hippocampus. The fHPSF #15 is the only fHPSF that connects with 22% of FPN.

The brain-wide FC profiles strongly depend on the selection of hippocampal seeds. The selection of seed regions that are functionally heterogeneous will cause reduced detection sensitivity and strength of FC. To understand how different hippocampal segmentations affect the spatial coverage of significant cotical-hippocampal subfield FC, Figure 6a and 6c demonstrate the hippocampal coverage of significant iFC associated with aHPSF and fHPSF respectively. For each cortical vertex, the hippocampal coverage of iFC is defined by the volumetric ratio of the significantly-connected subfields to the whole hippocampus. The cortical patterns associated with aHPSF and fHPSF appear to be similar but the values of the maps are different. To quantify the similarity, we generated the overlapping map (see Figure 6e) with the coverage threshold of 0. The green color encoded the overlapped areas which connected with at least one of the aHPSFs and fHPSFs. The blue and red colors encoded the areas which connected only with aHPSF and fHPSF, respectively. The corresponding dice coefficient is 0.84. Figure 6f showed that the overlapped area covered a majority of cortical surface, including 88% of visual network, 80% of sensorimotor network, 59% of DMN and 50% of frontoparietal network. On the other hand, the non-overlapped area did not show clear preferential FC with any functional network. Despite the similar pattern, nevertheless, the values of hippocampal coverages associated with aHPSF and fHSPF are apparently different. The radar charts in Figure 6b and 6d show the averaged hippocampal coverage within each functional network. The hippocampal coverages of fHPSF are generally higher than aHPSF, especially for visual network, sensorimotor network and DMN.

**Figure 6:**
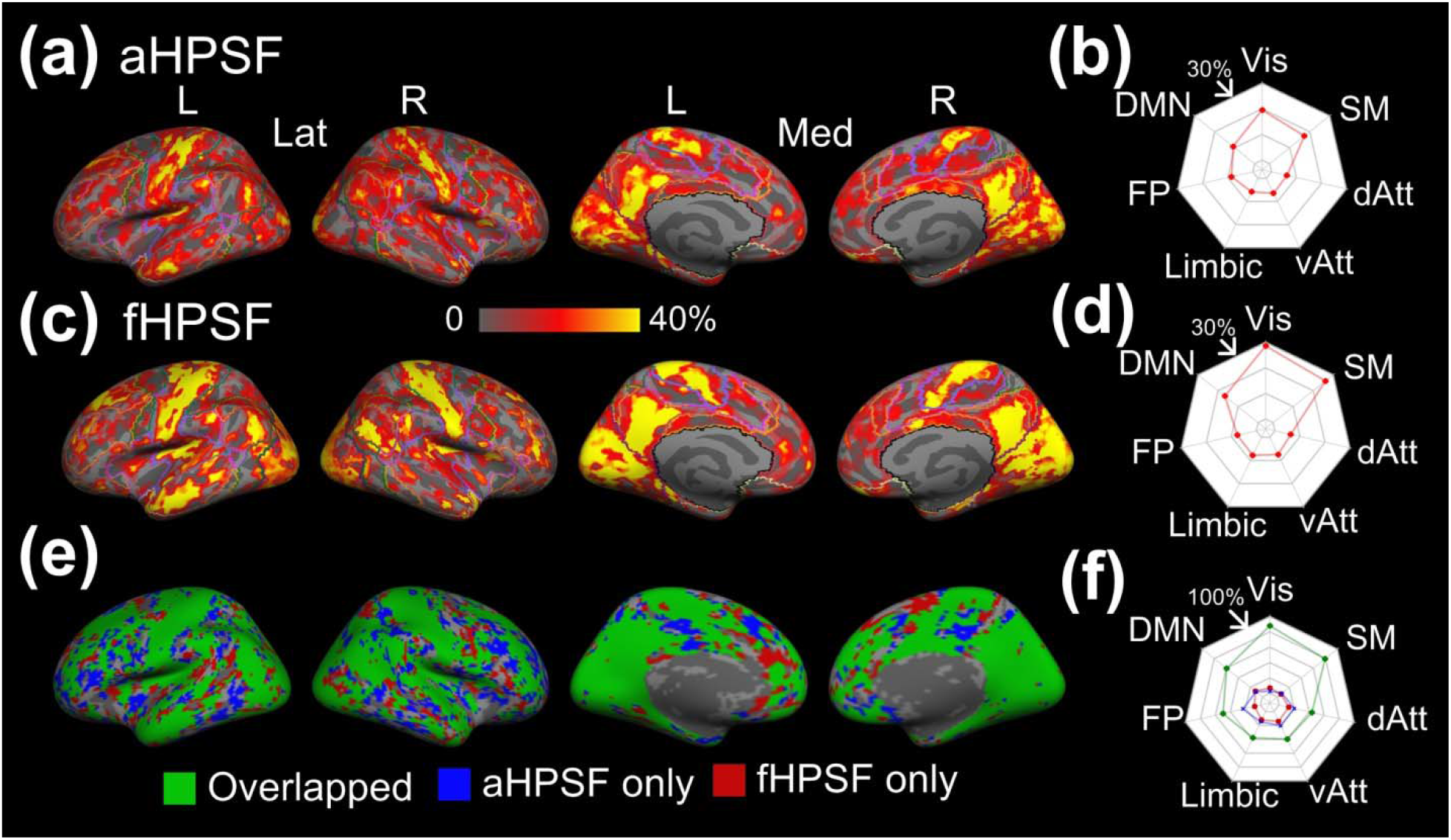
Hippocampal coverage of iFC associated with aHPSF and fHPSF. (a) The cortical map of hippocampal coverage associated with aHPSF. The contours of functional networks are outlined. (b) The radar chart of averaged hippocampal coverage within each functional network associated with aHPSF. (c) The cortical map of hippocampal coverage associated with fHPSF. (d) The radar chart of averaged hippocampal coverage within each functional network associated with fHPSF. (e) The overlap between the binarized cortical maps in (a) and in (b). The overlapping area is highlighted in green. The area which connects only with aHPSF is highlighted in blue; fHPSF in red. (f) The percentages of the overlapping, ‘aHPSF only’ and ‘fHPSF only’ area within each functional network. Abbreviations: L – left; R – right; Lat – lateral view; Med – medial view; Vis – visual network; SM – sensorimotor network; dAtt – dorsal attention network; vAtt – ventral attention network; Limbic – limbic network; FP – frontoparietal network; DMN – default-mode network.

## DISCUSSION

To minimize spurious functional similarity between hippocampal subfields, this study utilized an ultrahigh-resolution fMRI sequence (bPRISM) to achieve a submillimeter isotropic resolution, TR of 2s, and whole-brain coverage (Chang and Lin, 2018). With submillimeter fMRI, we were not only able to subdivide the hippocampus into lamellar structures such as subiculum, CA1, CA3, but also subdivided the lamellar structures into head and body portions. Additionally, we performed a fine-grained functional segmentation within the hippocampus using a data-driven approach. To the best of our knowledge, this is the first study to 1) delineate the hippocampal subfields functionally using ultrahigh-resolution fMRI, and to 2) compare the brain-wide FC associated with fine-grained aHPSFs and fHPSFs. Our analysis provided novel insights into the functional organization of hippocampus and the respective brain-wide networks.

### Functional Characteristics within Hippocampus

The fine-grained functional segmentation shown in Figure 2b demonstrated a spatial arrangement along hippocampal longitudinal axis, that largely reflected longitudinal functional specialization (Poppenk et al., 2013, Strange et al., 2014) without lamellar structures comparable to aHPSFs. As ultrahigh-resolution images minimize the signal mixture between subfields (as shown in Figure S4), our results mitigate the concern of partial volume effects on functional segmentation, as well as support the statement that anatomically-defined subfields do not represent the functional units of hippocampus at rest.

A recent study divided post-mortem human hippocampus into three portions (head, body and tail) and found that adjacent, but not distant, portions of each subfield were anatomically connected (Beaujoin et al., 2018). This finding dovetails with the result that nearly all the functional segments did not extend longitudinally from hippocampal head to tail. Outsides of the hippocampus, the afferent input to the entorhinal cortex have been reported to be organized in an anterior-to-posterior fashion (Kahn et al., 2008, Maass et al., 2015), which is largely preserved in its connections to the hippocampus (Maass et al., 2015). That is, anterior entorhinal cortices connect with anterior hippocampus and posterior entorhinal cortices connect with posterior hippocampus. These findings are consistent with our segmentation results as the longitudinally-arranged input pathways to the hippocampus corresponded with the longitudinally-arranged fHPSFs. The spatial arrangement of fHPSFs also clarified why the adjacent portions within or across aHPSFs were correlated, but distant portions were not (Dalton et al., 2019b), as most of the fHPSFs formed a cluster in proximity.

Although Dalton et al. (Dalton et al., 2019b) reported that adjacent, but not distal, segments within each aHPSF were correlated, CA1 was an exception. Interestingly, a similar phenomenon was observed in the current study. FHPSF #3 comprised ~56% of CA1 head and 41% of CA1 body. As shown in Figure 2b, the functional segment expanded from the anterior to posterior hippocampus. None of the other fHPSFs exhibited such a longitudinal expansion in shape. Hence, the significant iFC observed between distal portions of CA1 coincided with the functional homogeneity within the corresponding fHPSF. Whether it suggests that the longitudinal neuronal tracts exist exceptionally within CA1, or the distal portions within CA1 are connected polysynaptically, remains an open question.

The longitudinal fHPSFs parcellation observed in the current study is generally consistent with a recent report of functional segmentation (Plachti et al., 2019). However, the functional segmentation results in that study could not provide information about functional lateralization because the segmentations of left and right hippocampus were performed separately (Plachti et al., 2019). In contrast, we performed functional segmentation using left and right hippocampi jointly. Our results showed 13 out of 20 fHPSFs were unilateral, suggesting the coexistence of the unilateral and bilateral functional subfields. These results of hippocampal functional lateralization are in concordant with a number of prior studies. For example, left hippocampus is associated with more verbal memory processes, while right hippocampus with more spatially-dependent processes (Reuter-Lorenz et al., 2000, Burgess et al., 2002, Greve et al., 2011). Additionally, the CA1-like subregion in fHPSF was not observed in the earlier reports (Blessing et al., 2016, Plachti et al., 2019, Zhong et al., 2019). Many factors may contribute to this difference, including the spatial resolution of the fMRI data, the used de-noising approaches, and the employed clustering methods. Since one of our goals is to interrogate the respective brain-wide functional networks, the optimization of the functional segmentation was beyond the scope of this study. However, future investigation of the reliability and stability among functional segmentation approaches is an important area of inquiry.

### Functional Characteristics Related to Resting-State Functional Networks

Of the few studies examining FC in different portions of a hippocampal subfield (Libby et al., 2012, Maass et al., 2015, Dalton et al., 2019a, Dalton et al., 2019b), they investigated FCs at different portions of a subfield with the surrounding cortical areas, including posterior parahippocampal cortex (PHC) and perirhinal cortex (PRC). The features of brain-wide coverage and ultrahigh resolution in this study shifted the investigation of cortical-hippocampal subfield interactions from a regional level to a network level.

The functional segmentation of hippocampus provided more insights into the cortio-hippocampal subfield connectivity. The brain-wide iFC profiles across aHPSFs had been reported to be similar (Lacy and Stark, 2012, Shah et al., 2018, Vos de Wael et al., 2018). With the fine-grained functional segmentation of hippocampus, a number of fHPSFs showed the specificity of iFC with some of the functional networks, including visual, sensorimotor and default-mode networks. The functional-network specificity was not obvious with aHPSFs.

Our results demonstrate that visual and sensorimotor networks preferentially interact with different portions of CA1, CA3 and CA4/DG, as shown in Figure 5a. The visual and sensorimotor networks connect with the CA1 head more than CA1 body. On the contrary, the two functional networks show strong connections with CA3 body and CA4/DG body rather than the heads of CA3 and CA4/DG. As shown via histological studies, CA1 receives projections from inferior visual system and from V4 through parahippocampal and entorhinal cortices (Aggleton, 2012). Moreover, the ventral CA1 (corresponding to human anterior CA1) has direct projection to the olfactory bulb and several other primary olfactory cortical areas in rats (Cenquizca and Swanson, 2007) and monkeys (Roberts et al., 2007), lending support to the result that anterior CA1 connects with primary sensory cortices. In contrast, dorsal CA3 and CA4/DG are involved in the dorsal hippocampal network which is known to mediate cognitive process such as learning, memory, and exploration (Fanselow and Dong, 2010). Histological studies indicate CA3 and CA4/DG receive projections from superior visual system (Aggleton, 2012, Duvernoy et al., 2013). Our results of aHPSFs suggest that visual networks, as well as sensorimotor networks, connect with anterior CA1, posterior CA3 and CA4/DG. Consistent with our results of fHPSF, both the anterior and posterior subfields connect with visual and sensorimotor networks, but the posterior subfields are more dominant.

For DMN, the posterior pre-subiculum showed the most extensive FC. As documented in several recent subfield studies (de Flores et al., 2017, Vos de Wael et al., 2018), the subiculum shows more FC with DMN than CA1, CA3 and CA4/DG. Considering the close proximity between presubiculum and subiculum, the difference of spatial resolution and coregistration method approach may contribute to the discrepancy. For fHPSF, 5 out of the 6 subfileds that show extensive FC with DMN (≥ 20%) encompass body portions, suggesting that the posterior subfields show more extensive FC with DMN than anterior subfields. This is largely consistent with previous reports showing that posterior hippocampus is involved in the posterior medial (PM) system encompassing the dorsolateral/parietal components of DMN (Ranganath and Ritchey, 2012, Poppenk et al., 2013).

The FPN demonstrated smaller spatial extent of significant FC with hippocampal subfields compared to visual network, sensorimotor network and DMN in this study. Similar trend was also reported with respect to subiculum, combined subfield CA1-3 and CA4/DG (Vos de Wael et al., 2018). Since FPN serves as a flexible hub of cognitive control (Zanto and Gazzaley, 2013), weaker FC with hippocampal subfields during resting state is biologically plausible. Moreover, our results demonstrated that the lateral prefrontal and posterior parietal cortices in FPN were anticorrelated with the head portion of CA1 (see Figure 3a). Such an anticorrelation of FPN with anterior hippocampus was also observed in a recent study (Zhong et al., 2019). Although the anticorrelation between DMN and FPN had been reported to reflect the competitive balance between internally self-referential processing and externally oriented cognitive processing (Uddin et al., 2009), the anticorrelation between FPN and hippocampal subfields are less explored. More experiments are needed to clarify its biological role. Our group has an ongoing research to investigate the functional role of FPN-hippocampal subfield anticorrelation by different task challenges.

### Selection of Hippocampal Segmentation – Anatomical or Functional?

The iFC profiles of hippocampus with respect to the RSNs are determined by the selection of hippocampal segmentation. While results of the current study suggest that using fHPSFs as the seeds may have the advantage of higher FC strength (see Figure 6), functional segmentation of the hippocampus requires a certain level of statistical power and is usually more complicated, especially for clinical applications. Other than the difference of FC strength, however, the cortical regions that are connected with any of aHPSF are highly overlapped with those of fHPSF (see Figure 6e). In other words, using aHPSF for iFC analysis may compromise the strength of FC, but not the presence of FC.

An enhanced understanding of functional segmentation of the hippocampus and the respective cortical networks holds a great promise for clinical applications. The hippocampus is the earliest and most severely affected structure in neurodegenerative disorders such as Alzheimer’s disease (AD) (Jack et al., 2013). Based on the network degeneration hypothesis (Seeley et al., 2009), the pathological process of AD could occur selectively within the fHPSFs and the respective functional networks. Therefore, the findings in this study provide fundamental information to potentially predict pathological effects. Moreover, the investigation of the functional networks associated with the fHPSFs may potentially enhance the sensitivity of AD biomarkers, as well as biomarkers for a range of other disorder shown to have structural alterations in hippocampal subfields including autism, Alzheimer’s disease, schizophrenia, and bipolar disorder, as network-based approaches take into account disease-related alterations in brain-wide interconnections rather than measuring the regional changes of hippocampus.

### Limitations

Some limitations should be considered in the current study. First, all of the function images were acquired using a 7T scanner. Therefore, strong susceptibility artifacts may affect the signal quality around the air-tissue interface. In this study, the structures around the anterior and inferior temporal lobes such as the entorhinal cortex, perirhinal cortex, fusiform gyrus, temporal pole experienced severe signal loss (see Figure S5 in the Supplementary materials for details). The inferior part of the hippocampus was also be affected. Although we analyzed the temporal signal-to-noise ratio (tSNR) of each hippocampal subfield and regressed out the tSNR variation for the group analysis, the reduction of iFC sensitivity due to signal loss is inevitable. As such, the results related to those regions may not be as reliable.

Secondly, the anatomical segmentation in this study was performed based on the T1-weighted image with 0.94 mm isotropic resolution. Since FSL stores the discrete segmentation volume at 0.33 mm resolution, it is unclear whether 0.94 mm resolution is sufficient for accurate hippocampal segmentation. To test this issue, we acquired 16 datasets of T2-weight images with 0.6-mm isotropic resolution and T1-weighted images with 1-mm isotropic resolution. The hippocampal segmentation was performed using T1-weighted and T2-weighted images separately. The similarity of the hippocampal segmentations was measured by dice coefficient. As shown in Figure S7 (see Supplementary materials), the two segmentation results were highly similar with an overall dice coefficient of 0.99. This finding suggests that the quality of hippocampal segmentation with 0.94-mm resolution is relatively reliable.

### Technical Notes

Of note, it is not our intention to claim that the proposed functional organization of hippocampus is the absolutely accurate model. Rather, this study demonstrates efforts to probe the functional organization of hippocampus by minimizing the biases from insufficient spatial resolution and a priori subfield delineation. Imaging with lower spatial resolution may not only compromise the boundary details, but also reduce the statistical power for multiple comparisons. For studies that focus on subtle changes of subfield boundaries, such as the aging effect on the functional organization of hippocampus, ultrahigh-resolution functional imaging provides unique advantages.

## CONCLUSIONS

This study developed and utilized fMRI data acquisition with submillimeter isotropic resolution, whole-cerebrum coverage, and TR of 2s to 1) investigate whether hippocampal functional segmentation with ultrahigh-resolution data demonstrate similar anatomical, lamellar structures in the hippocampus, and 2) define and compare the brain-wide FC associated with fine-grained aHPSFs and fHPSFs. No clear lamellar structures were observed in our results of functional segmentation, suggesting that anatomically-defined hippocampal subfields do not represent the functional units of hippocampus at rest. The spatial arrangement of fHPSFs was primarily along the longitudinal axis. While most of the aHPSFs are quite different from the fHPSFs, CA1 was the unique exception. For brain-wide functional networks, we found that the fHPSFs rather than aHPSFs revealed specific FC with some of the functional networks. The visual and sensorimotor networks preferentially interact with different portions of CA1, CA3 and CA4/DG. Visual, sensorimotor, and default-mode networks showed more prominent FC with the body portion of hippocampus than with the head portion. Moreover, lateral prefrontal and posterior parietal cortices in FPN were shown to be anticorrelated with the head portion of CA1. Taken together, our results of fine-grained functional segmentation and the respective functional networks provide valuable insights into the applications of neurodegenerative diseases.

## ACKNOWLEGEMENTS

This work was supported in part by NIH grants, R21NS095027, R21AG060324 and U01MH110274.

## Notes

### Competing Interest Statement

The authors have declared no competing interest.

