## Supplementary materials for "Brain-wide functional networks associated with anatomically- and functionally-defined hippocampal subfields using ultrahigh-resolution fMRI"

**Blipped Partition-encoded Simultaneous Multi-Slab (bPRISM)**

In recent years, simultaneous multislice imaging (SMS) has become the imaging method of choice for fMRI due to its ability to accelerate image acquisition along the slice direction (Setsompop et al., 2012, Setsompop et al., 2013, Barth et al., 2016, Setsompop et al., 2016). However, the resolution of SMS along the slice direction is limited by the physical constraints of slice excitation. Thin slice excitation is challenging due to the requirements of a high gradient, well-defined RF excitation profile and lengthened RF pulse duration. Alternatively, the simultaneous multislab (SMSlab) approaches achieve a thin slice thickness without the aforementioned physical constraints through performing Fourier encoding within a slab (Chen and Feinberg, 2013, Engstrom et al., 2015, Bruce et al., 2017). Our group recently proposed the full-sampling principles of SMSlab in 3D k-space purely by gradient encoding, consisting of both inter-slab and intra-slab encoding (Chang et al., 2018). Based on the fully-sampled pattern of SMSlab, we defined the acceleration pattern using blipped CAIPI technique, which we termed blipped PRISM (bPRISM). The bPRISM sequences achieve submillimeter isotropic resolution, whole-cerebrum coverage and temporal resolution of 2 seconds, which is critically important for the study of cortical-hippocampal subfield functional connectivity.

**Simultaneous Multi-slab Excitation**

To perform simultaneous multi-slab excitation, conventional slice-selective RF pulses were frequency modulated and summed. To minimize the peak RF amplitude, the optimized set of excitation phases across slices was applied (Wong, 2012). The RF pulses were designed based on the shinnar-Le Roux (SLR) algorithm (Pauly et al., 1991). The RF pulse with small flip angle (≤ 60°) and excitation pulse with larger flip angle (60° < flip angle ≤ 90°) were designed separately using Spectral-Spatial RF Pulse Design for MRI and MRSI MATLAB Package (<http://rsl.stanford.edu/research/software.html>) (Larson et al., 2008).

**MR Acquisitions**

The reference and calibration scans are combined in the bPRISM sequences. Conventionally, the reference scan for SMS reconstruction and the calibration scan for inplane unaliasing were acquired separately before the acceleration scan. Consequently, the consistency between reference and calibration scan may become an issue. The long combined acquisition time also makes the image prone to degradation induced by head movement and physiological fluctuation. This study shortened the acquisition time of reference and calibration scans by acquiring a fully-sampled reference scan using segmented-EPI method. The calibration scan was discarded because the fully-sampled reference scan encompasses the auto-calibration scan (ACS) data already. Assuming the in-plane acceleration rate is 2, the acquisition time will be shortened by 33%.

**MR Reconstructions**

The slice-GRAPPA and inplane-GRAPPA are typically used sequentially for SMS reconstruction. As a result, the reconstruction error in the first reconstruction step propagates to, and is enhanced by, the next step. In contrast, this study performed the SMS reconstruction in one step as shown in Figure S1, thereby lowering the g-factor penalty (Breuer et al., 2009). The final step is to perform spatial decoding along the partition encoding (PAE) direction. The Fourier bases in the spatial decoding matrix are mutually orthogonal, and therefore, the through-plane information within a slab can be unambiguously reconstructed.

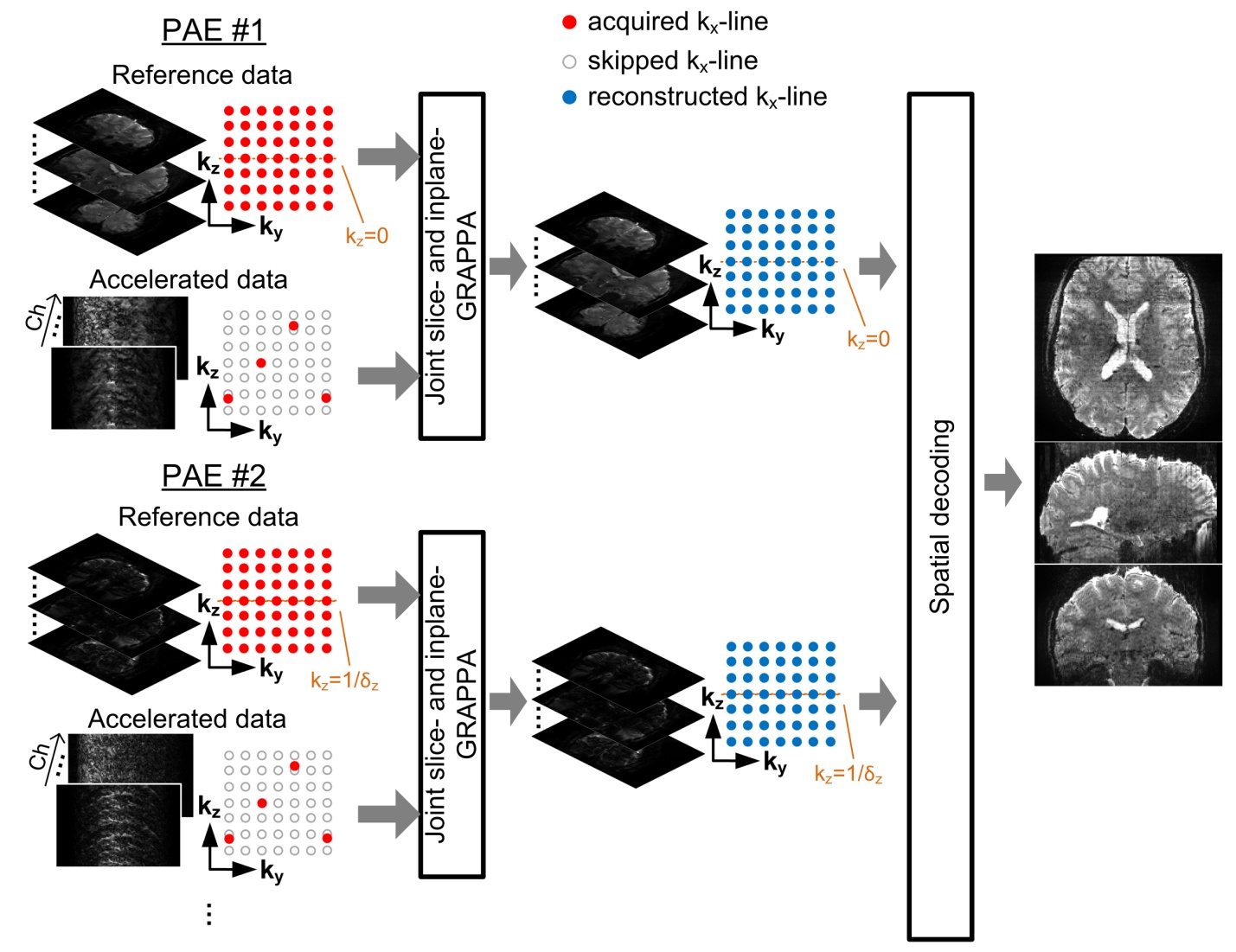

Figure S1: The image reconstruction procedure. The kernels of joint slice and in-plane GRAPPA are estimated for each partition encoding. The filled red, unfilled and filled blue circles represent the sampled, skipped and reconstructed k_x_ lines respectively. δ_z_ denotes the slab thickness. Abbreviations – PAE: partition encoding, Ch: channel.

**Sensitivity analysis of bPRISM**

Compared to SMS, the sensitivity of bPRISM could be affected by two major factors: the reduced time interval between two contiguous excitation (TR_ex_), resulting in a lower steady-state signal, and a slab n_p_ times thicker than a slice, leading to a SNR advantage of $\sqrt{\text{n}_{\text{p}}}$. The sensitivity *S* is defined as

$\text{S=}\sqrt{\text{n}_{\text{p}}}\cdot\text{M}_{z}={\sqrt{\text{n}_{\text{p}}}\text{∙}\text{M}}_{0}sin\theta\left( 1-exp\left( -\text{TR}_{\text{ex}}/T_{1} \right) \right)/\left( 1-cos\theta\cdot exp\left( -\text{TR}_{\text{ex}}/T_{1} \right) \right)$,	(1)

where $\theta$, M_z_, and M_0_ are the flip angle, steady-state longitudinal magnetization and longitudinal magnetization at equilibrium, respectively. For simplicity, M_0_ was set as 1 and a T_1_ of 1900 ms was employed for simulation, which is a typical T1 value of gray matter at 7T (Wright et al., 2008).

**Performance Comparison Between SMS and bPRISM Sequences**

Assuming the TR = (number of PAE)×TR_ex_ is a constant value, different number of PAE will lead to different sensitivity as calculated by Eq. (1). According to the theoretical sensitivity shown in Figure S2a, the bPRISM showed only ~5% increase compared to SMS for acceleration scans. However, bPRISM acquisitions not only increase the sensitivity of acceleration but also of reference scans. Figure S2b indicates that the sensitivity increase in the bPRISM sequences was >75% higher than SMS sequences. The high quality of reference images yielded the reconstruction kernel more accurate, thereby lowering the g-factor penalty. Figure S2c demonstrates the temporal SNR (tSNR) of SMS, bPRISM and the enhancement. The tSNR of bPRISM is profoundly higher than that of SMS. The averaged enhancement across the entire brain was 16.9%. The slab boundary effect in the tSNR map of bPRISM was not pronounced. The averaged tSNR of bPRISM at the edge slices was ~4% lower than the non-edged slices.

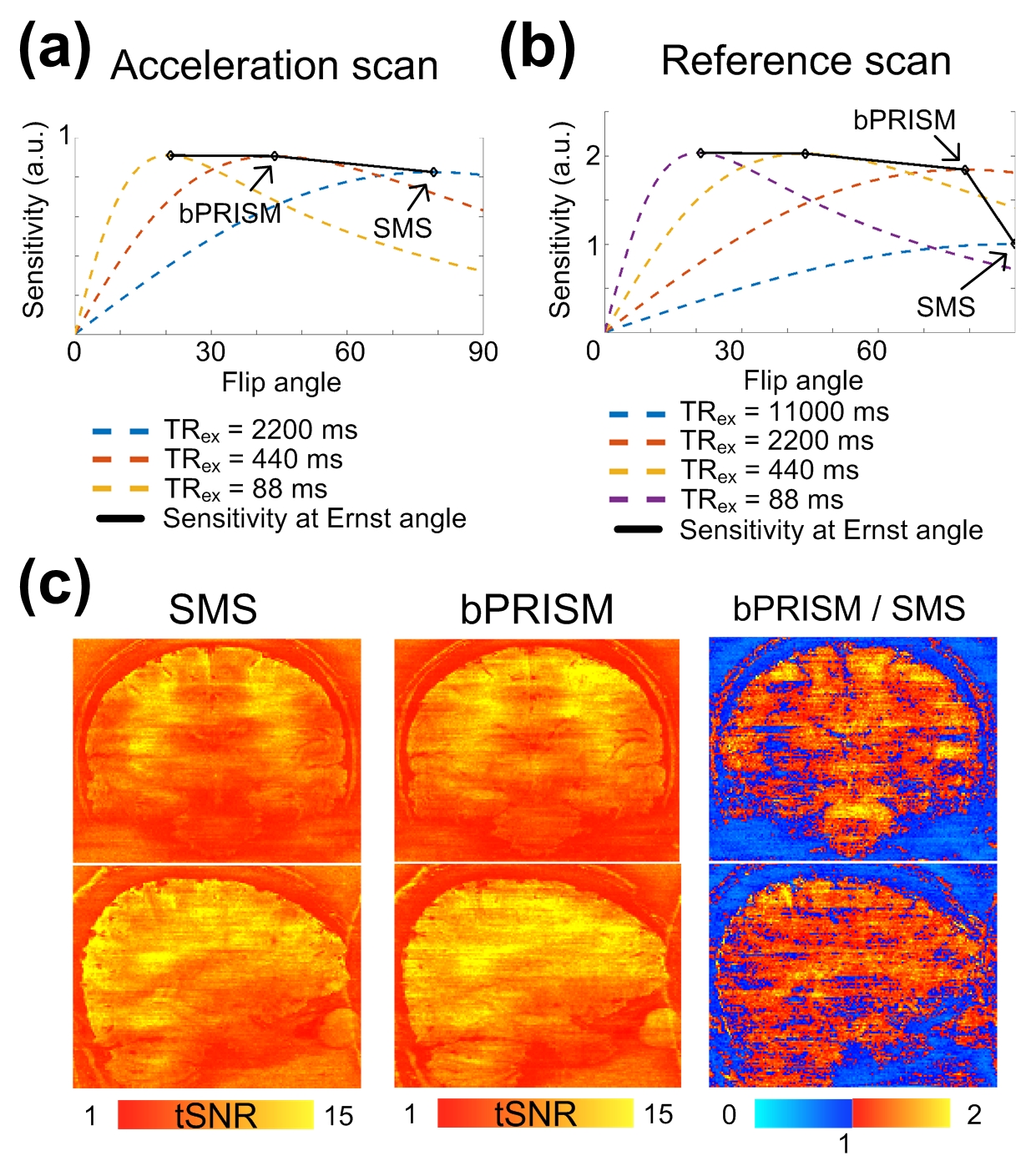

Figure S2. Performance comparison between SMS and bPRISM. (a) The sensitivity traces against flip angles for acceleration scan. The TR of acceleration scan is 2200 ms and is equal to (number of PAE) × TR_ex_. The traces associated with different TR_ex_ are color-encoded as indicated in the figure legend. (b) The sensitivity traces against flip angles for reference scan. The TR of reference scan is 11000 ms. The traces associated with different TR_ex_ are color-encoded as indicated in the figure legend. (c) The tSNR and the enhancement maps. The spatial resolution is 1 mm isotropic and the TR is 2.2s. The SMS factor is 5. Total number of axial slices is 125.

**Resting-State Functional Networks with Sub-Millimeter Isotropic Resolution**

Figure S3 shows the connectivity patterns of resting-state networks using submillimeter fMRI. The result was obtained from two 6-minute resting runs of single participant with 0.94-mm isotropic resolution. Minimum spatial smoothing of 0.1 mm was used. The majority of the connectivity patterns were confined in gray matter. Several typical patterns of resting-state networks were observed, such as visual medial network, visual occipital network, default mode network (DMN) and executive control network (ECN). The results suggest that single participant submillimeter fMRI using PRISM was sensitive enough to detect resting-state functional networks.

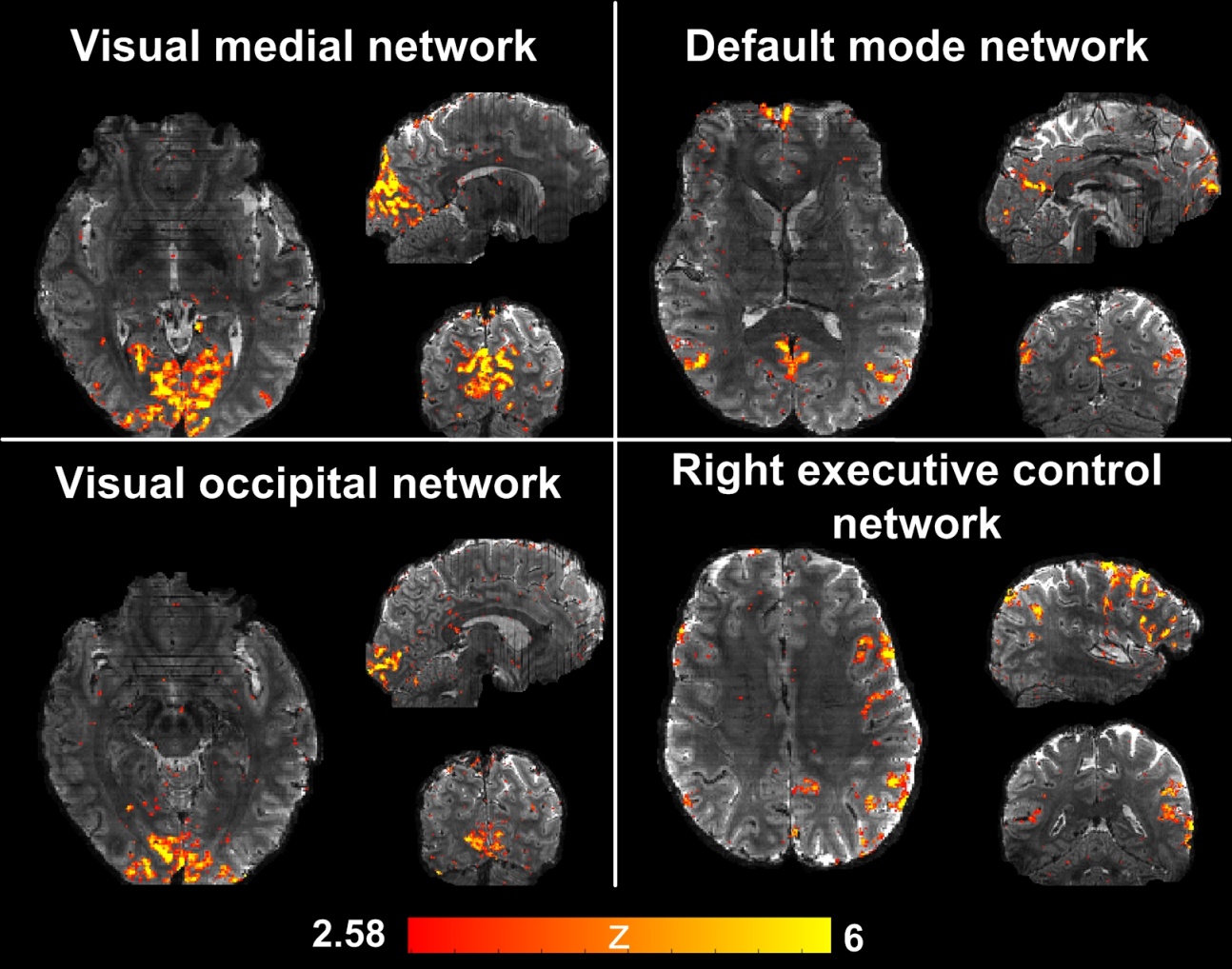

Figure S3. Resting-state functional networks obtained from a single subject using 0.94mm isotropic resolution. The names of the functional networks are shown as the panel title. The z statistics are color-encoded as indicated by the color bar and overlaid on the reference image.

**Partial Volume Effect on the Inter-Subfield Functional Correlations**

While the intrinsic functional connectivity (iFC) between whole hippocampus and the rest of the brain is well established, studies investigating the functional networks associated with the hippocampal subfields are relatively few mainly because the hippocampus curls into S-shaped structure in a small volume of 3-3.5 cm^3^ and the width of hippocampal subfield could be less than a millimeter. Hence, ultrahigh resolution functional imaging is preferable to alleviate the partial volume effect. Nevertheless, acquiring functional MRI (fMRI) with ultrahigh spatial isotropic resolution and whole-brain coverage within a 2-second time frame is technically challenging. Many research groups have studied cortical-hippocampal subfield interactions using the fMRI with the spatial resolution of ≥ 2 mm isotropic (de Flores et al., 2017, Li et al., 2018, Zhong et al., 2019). However, the corresponding partial volume effects on the inter-subfield functional correlations have not been systematically investigated.

This study performed simulations to investigate how spatial resolution affects functional correlations between hippocampal subfields. First, we acquired a T2-weighted image with the resolution of 0.6 mm isotropic using turbo-spin-echo (TSE) sequence and then performed the hippocampal segmentation using Freesurfer (Fischl, 2012, Iglesias et al., 2015), as shown in Figure S4a. Second, we deliberately selected 5 time courses with low correlations from *in vivo* datasets. The 5 time courses were used as the time courses of the hippocampal subfields, as shown in Figure S4b. Specifically, time series with 0.6-mm isotropic resolution were synthesized by adding the time courses onto the entire hippocampal subfields. The time courses had zero means and the standard derivation of each voxel was equal to 2% of the voxel intensity. The left-most column in Figure S4c shows the temporal average of the image series with 0.6 mm isotropic resolution and the corresponding correlation matrix. Images with 1-mm, 2-mm and 3-mm isotropic resolutions were obtained by downsampling the image series of 0.6-mm resolution. The downsampling was implemented by discarding the high-frequency portions in the k-space. Additionally, we simulated the effect of T2* decay along the phase-encoding dimension (superior-inferior direction) with echo-spacing of 1 ms and T2* of 32 ms.

The resulting correlation matrices in the 2^nd^, 3^rd^ and 4^th^ columns in Figure S4c suggests that large voxel size induced spurious correlations between hippocampal subfields. Figure S4d shows the means and mean square errors of different spatial resolutions. The trend of increased functional correlations indicates that lower spatial resolution will cause higher false positives of inter-subfield functional similarity.

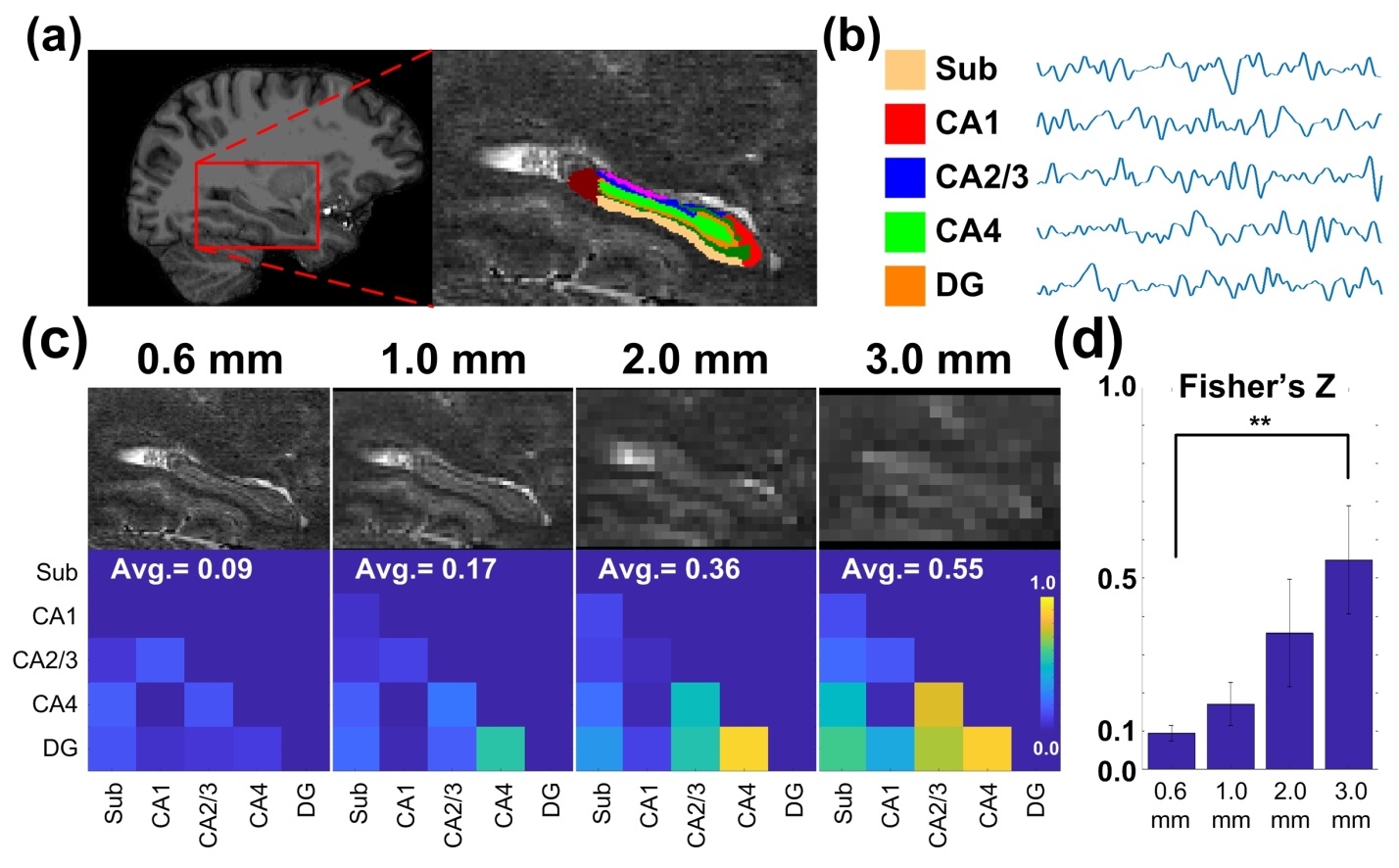

Figure S4: The simulated partial volume effects on inter-subfield functional correlations. (a) Hippocampus image with 0.6-mm isotropic resolution and the overlaid labels of hippocampal subfields. The right panel is the enlarged image from the red block of the left panel. (b) The simulated time courses of the hippocampal subfields. (c) The upper row shows the images of hippocampus with 0.6-mm, 1-mm, 2-mm, 3-mm isotropic resolutions and the bottom row showed the corresponding functional correlations between hippocampal subfields. (d) The means and mean square errors of inter-subfield functional correlations after the Fisher’s Z transformation. The whiskers represent the mean square errors. Statistical significance of the paired t-test is indicated as **: p < 0.01. Abbreviations: Sub – subiculum; CA – cornu ammonis; DG – dentate gyrus.

**Quantification of Signal Quality**

The signal loss due to the susceptibility effect was quantified by tSNR. The tSNR of each individual was calculated before any data processing, i.e. using raw data. The tSNR values on the native EPI space were warpped onto the individual T1 image before being re-sampled onto individual cortical surface. The re-sampling of tSNR values on EPI space captured only the values from the white matter surface to the middle depth of cerebral cortex in order to suppress the contamination from large vessels. The individual surface data were finally warpped onto the template surface and averaged as shown in Figure S5a. The signal quality around the inferior brain such as temporal lobe (5.4 < tSNR < 8.2), orbital frontal lobe (tSNR = 6.4) are relatively low.

The tSNR within individual hippocampus were warpped onto the template brain. The details of the coregistration are as described in the **Functional Segmentation of Hippocampus** section. The mean and standard deviation of tSNR within each hippocampal subfield were shown in Figure S5b. In general, the tSNRs in the hippocampal head are lower than the hippocampal body and tail.

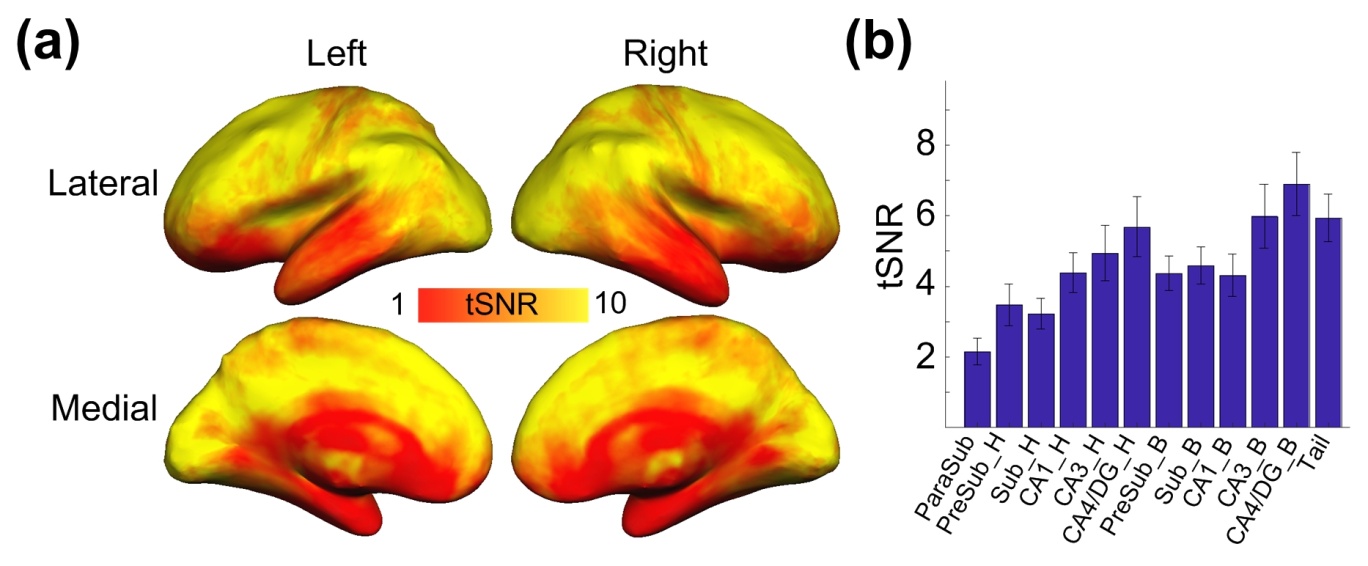

Figure S5: The group-averaged tSNR of raw data. (a) The tSNR on the cortical surface. The tSNR values are encoded as indicated by the colorbar. (b) The tSNR of each hippocampal subfield. Abbreviations: H – head; B – body; Sub – subiculum; CA – cornu ammonis; DG – dentate gyrus.

**Functional segmentation procedures**

The procedure of functional segmentation is shown in Figure S6. Although the transformation from individual EPI to the template space may cause spatial blurring among hippocampal time courses, such spatial blurring did not affect the final segmentation results directly because of the soft segmentation, i.e. k-means clustering. The k-means clustering method in this study utilized the probability maps of all the independent components as the feature vectors. The spatial blurring induced by the nonlinear coregistration altered the probabilities of the independent components, causing a shift in the space of feature vectors. If two adjacent voxels were functionally distinct, the distance between them in the vector space was long such that the relatively slight shift would not affect the result of clustering; if the two adjacent voxels were functionally close to each other, the impact of blurring on the clustering result becomes inconsequential in that the two voxels have been functionally similar anyway. Therefore, spatial blurring did not impact the final results of functional segmentation as directly as of functional connectivity.

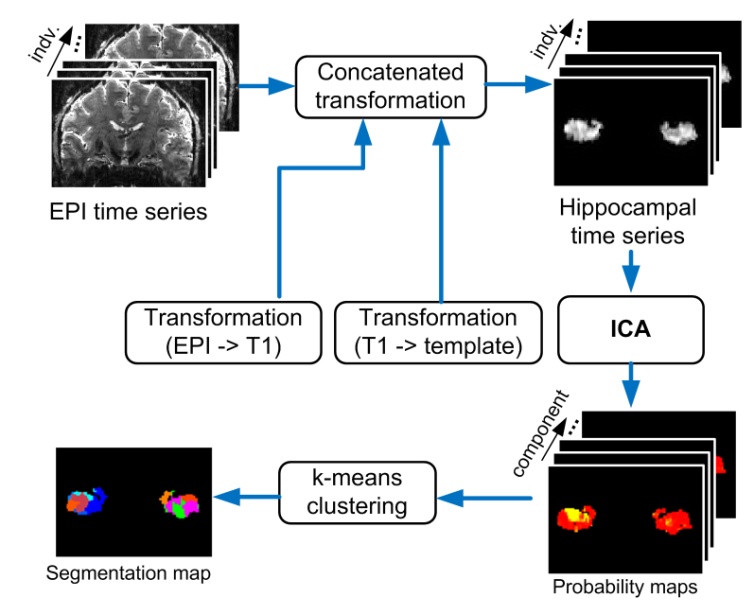

Figure S6: Functional segmentation procedures. Hippocampal time series of all subjects were transformed onto the template space. The coregistered hippocampal time series were passed to ICA and produced probability maps of all independent components. The k-means clustering method utilized the probability maps as the feature vector and generated the segmentation map. The initial cluster number was set manually and then determined automatically according to the elbow criterion of the cluster validity index.

**Reliability of Anatomical Hippocampal Segmentation**

The anatomical segmentation in this study was performed based on the T1-weighted image with 0.94 mm isotropic resolution. Since the FSL store the discrete segmentation volume at 0.33 mm resolution, it is not clear whether 0.94 mm resolution is sufficient for accurate hippocampal segmentation. To test this issue, we acquired 16 datasets of T2-weight images with 0.6-mm isotropic resolution and T1-weighted images with 1-mm isotropic resolution. The hippocampal segmentation was performed using T1-weighted and T2-weighted images separately.

MR images were acquired using a Siemens 7T scanner. High-resolution MPRAGE image was acquired using the imaging parameters as follows: TR/TE/TI = 2200/2.78/1050 ms, flip angle = 7°, partition thickness = 1 mm, image matrix = 256 × 240, 192 partitions, and FOV = 25.6 cm × 24.0 cm. Additionally, ultrahigh-resolution TSE image was also acquired for T2-weighted image. Due to the limited scan time, only an axial slab that encompassed the hippocampus was acquired. The imaging parameters are as follows: spatial resolution = 0.6 mm isotropic, TR/TE = 14870/67 ms, flip angle = 179°, slice thickness = 0.6 mm, image matrix = 304 × 288, 72 slices, and FOV = 18.2 cm × 17.26 cm, in-plane GRAPPA acceleration factor = 2, Turbo factor = 8, echo train per slice = 20.

Figure S7a demonstrated the anatomical segmentation maps of hippocampus from a representative participant. The middle and right panels showed the segmentation maps using 1-mm isotropic T1-weighted image and 0.6-mm isotropic T2-weighted image respectively. The two segmentation maps are similar qualitatively. The similarity was quantified by dice coefficient as shown in Figure S7b. Most of the dice coefficients are closed to 1 and the overall average is 0.99. This suggests the quality of hippocampal segmentation with 0.94-mm resolution is relatively reliable.

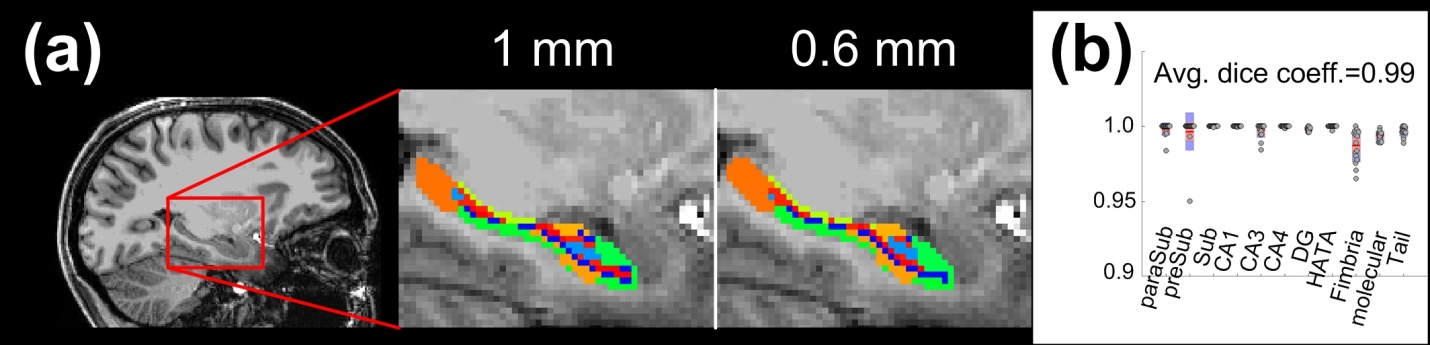
Figure S7: The anatomical segmentation using 1-mm isotropic and 0.6-mm isotropic T1-weighted images. (a) The background images in the middle and right panels were enlarged from the red box in the left panel. The spatial resolutions of the background images are both 1-mm isotropic. The overlaid segmentation map in the middle and right panels were derived from 1-mm isotropic T1-weighted image and 0.6-mm isotropic T2-weighted image respectively. Different colors encode different subdivisions of hippocampus. (b) The dice coefficients of subdivisions and individuals between 1-mm and 0.6-mm results. Each dot represents the dice coefficient of one subdivision of one individual. The dice coefficient averaged across subdivisions and individuals is 0.99. Abbreviations: paraSub – parasubiculum; preSub – presubiculum; Sub – subiculum; CA – cornu ammonis; DG – dentate gyrus; HATA – hippocampus-amygdala-transition-area; Molecular – molecular layer; Tail – hippocampal tail.

### RFERENCES

Barth M, Breuer F, Koopmans PJ, Norris DG, Poser BA (2016) Simultaneous multislice (SMS) imaging techniques. Magn Reson Med 75:63-81.

Breuer FA, Kannengiesser SA, Blaimer M, Seiberlich N, Jakob PM, Griswold MA (2009) General formulation for quantitative G-factor calculation in GRAPPA reconstructions. Magn Reson Med 62:739-746.

Bruce IP, Chang HC, Petty C, Chen NK, Song AW (2017) 3D-MB-MUSE: A robust 3D multi-slab, multi-band and multi-shot reconstruction approach for ultrahigh resolution diffusion MRI. Neuroimage 159:46-56.

Chang WT, Puspitasari F, Garcia-Miralles M, Yeow LY, Tay HC, Koh KB, Tan LJ, Pouladi MA, Chuang KH (2018) Connectomic imaging reveals Huntington-related pathological and pharmaceutical effects in a mouse model. NMR Biomed e4007.

Chen L, Feinberg D (2013) Simultaneous Multi-Slab Echo Volume Imaging: comparison in sub-second fMRI. In: Proceedings of the ISMRM Salt Lake City, United States.

de Flores R, Mutlu J, Bejanin A, Gonneaud J, Landeau B, Tomadesso C, Mezenge F, de La Sayette V, Eustache F, Chetelat G (2017) Intrinsic connectivity of hippocampal subfields in normal elderly and mild cognitive impairment patients. Hum Brain Mapp 38:4922-4932.

Engstrom M, Martensson M, Avventi E, Skare S (2015) On the signal-to-noise ratio efficiency and slab-banding artifacts in three-dimensional multislab diffusion-weighted echo-planar imaging. Magn Reson Med 73:718-725.

Fischl B (2012) FreeSurfer. Neuroimage 62:774-781.

Iglesias JE, Augustinack JC, Nguyen K, Player CM, Player A, Wright M, Roy N, Frosch MP, McKee AC, Wald LL, Fischl B, Van Leemput K, Alzheimer's Disease Neuroimaging I (2015) A computational atlas of the hippocampal formation using ex vivo, ultra-high resolution MRI: Application to adaptive segmentation of in vivo MRI. Neuroimage 115:117-137.

Larson PE, Kerr AB, Chen AP, Lustig MS, Zierhut ML, Hu S, Cunningham CH, Pauly JM, Kurhanewicz J, Vigneron DB (2008) Multiband excitation pulses for hyperpolarized 13C dynamic chemical-shift imaging. J Magn Reson 194:121-127.

Li H, Jia X, Qi Z, Fan X, Ma T, Pang R, Ni H, Li CR, Lu J, Li K (2018) Disrupted Functional Connectivity of Cornu Ammonis Subregions in Amnestic Mild Cognitive Impairment: A Longitudinal Resting-State fMRI Study. Front Hum Neurosci 12:413.

Pauly J, Le Roux P, Nishimura D, Macovski A (1991) Parameter relations for the Shinnar-Le Roux selective excitation pulse design algorithm [NMR imaging]. IEEE Trans Med Imaging 10:53-65.

Setsompop K, Feinberg DA, Polimeni JR (2016) Rapid brain MRI acquisition techniques at ultra-high fields. NMR Biomed 29:1198-1221.

Setsompop K, Gagoski BA, Polimeni JR, Witzel T, Wedeen VJ, Wald LL (2012) Blipped-controlled aliasing in parallel imaging for simultaneous multislice echo planar imaging with reduced g-factor penalty. Magn Reson Med 67:1210-1224.

Setsompop K, Kimmlingen R, Eberlein E, Witzel T, Cohen-Adad J, McNab JA, Keil B, Tisdall MD, Hoecht P, Dietz P, Cauley SF, Tountcheva V, Matschl V, Lenz VH, Heberlein K, Potthast A, Thein H, Van Horn J, Toga A, Schmitt F, Lehne D, Rosen BR, Wedeen V, Wald LL (2013) Pushing the limits of in vivo diffusion MRI for the Human Connectome Project. Neuroimage 80:220-233.

Wong E (2012) Optimized Phase Schedules for Minimizing Peak RF Power in Simultaneous Multi-Slice RF Excitation Pulses. ISMRM 20th Annual Meeting & Exhibition. Melbourne, Australia.

Wright PJ, Mougin OE, Totman JJ, Peters AM, Brookes MJ, Coxon R, Morris PE, Clemence M, Francis ST, Bowtell RW, Gowland PA (2008) Water proton T1 measurements in brain tissue at 7, 3, and 1.5 T using IR-EPI, IR-TSE, and MPRAGE: results and optimization. MAGMA 21:121-130.

Zhong Q, Xu H, Qin J, Zeng LL, Hu D, Shen H (2019) Functional parcellation of the hippocampus from resting-state dynamic functional connectivity. Brain Res 1715:165-175.
